# Uncovering Hidden Cancer Self-Dependencies through Analysis of shRNA-Level Dependency Scores

**DOI:** 10.1101/2023.03.23.533901

**Authors:** Zohreh Toghrayee, Hesam Montazeri

## Abstract

Large-scale short hairpin RNA (shRNA) screens on well-characterized human cancer cell lines have been widely used to identify novel cancer dependencies. However, the off-target effects of shRNA reagents pose a significant challenge in the analysis of these screens. To mitigate these off-target effects, various approaches have been proposed that aggregate different shRNA viability scores targeting a gene into a single gene-level viability score. Most computational methods for discovering cancer dependencies rely on these gene-level scores. In this paper, we propose a computational method, named NBDep, to find cancer self-dependencies by directly analyzing shRNA-level dependency scores instead of gene-level scores. The NBDep algorithm begins by removing known batch effects of the shRNAs and selecting a subset of concordant shRNAs for each gene. It then uses negative binomial random effects models to statistically assess the dependency between genetic alterations and the viabilities of cell lines by incorporating all shRNA dependency scores of each gene into the model. We applied NBDep to the shRNA dependency scores available at Project DRIVE, which covers 26 different types of cancer. The proposed method identified more well-known and putative cancer genes compared to alternative gene-level approaches in pan-cancer and cancer-specific analyses. Additionally, we demonstrated that NBDep controls type-I error and outperforms statistical tests based on gene-level scores in simulation studies.

**Author Summary:** Large-scale shRNA screening is increasingly being used in cancer genomics to discover genes involved in cancer by analyzing the viabilities of cell lines upon knocking down a gene using a pool of short hairpin RNAs (shRNA). However, off-target effects, which result from the knockdown of unintended genes, are a major issue in RNAi screening. To address this issue, various computational methods have been developed to aggregate shRNA viability scores into gene-level dependency scores. In this paper, we propose a method called NBDep to identify cancer gene drivers that directly addresses the challenge of off-target effects at the shRNA level. NBDep identifies cancer gene drivers in three classes: amplification, missense, and non-missense alterations. In this method, we first remove known batch effects, select a subset of the most consistent shRNAs of each gene, and then perform a negative binomial mixed-effect model. The NBDep method not only identifies well-recognized and novel cancer driver genes but also has more statistical power than gene-level-score methods while controlling type-error. In summary, NBDep presents a new technique for analyzing shRNA screens and has the potential to uncover previously unknown cancer dependencies.

## 1 Introduction

Large-scale CRISPR-Cas9 and RNAi screens have been increasingly used in cancer research to identify novel cancer vulnerabilities and therapeutic choices. While CRISPR-Cas9 technology can be used to perform knockout of gene function at the DNA level through multiple single-guide RNAs, RNAi screens knockdown genes at the mRNA level using a pool of short hairpin RNAs (shRNA) [1-6]. Despite the potential benefits of these screens in identifying cancer dependencies, a major challenge in the analysis of pooled shRNA screens is to account for the off-target effects of reagents[7, 8]. Various computational approaches have been proposed to mitigate the impact of off-target effects in the analysis of shRNA screens [9-12]. A common theme among all the previous approaches for identifying cancer dependencies is to first aggregate individual reagent effects into a single gene-level score using various computational tools such as RSA [12], ATARIS [9], and DEMETER [13]. Gene-level scores are then used to compare subjects with highly diverse molecular profiles in order to infer cancer dependencies.

The RSA method employs a probabilistic approach to calculate absolute gene-level viability scores from multiple siRNAs targeting a specific gene. It evaluates whether the siRNAs targeting the gene are unusually top-ranked among all siRNAs in the screen [12]. The ATARiS method provides relative gene-level dependency values by only incorporating a subset of RNAi reagents whose phenotypic effects are concordant across multiple samples [9]. DEMETER is another computational framework for estimating relative gene-level scores using multiple shRNA effects, assuming that each observed shRNA value is a linear combination of the corresponding gene-level effect and the batch effect of the corresponding seed sequence [13]. The gene-level effects are estimated using a stochastic gradient descent algorithm to minimize a regularized objective function. DEMETER2 extends the original DEMETER by using a hierarchical model for the gene and seed effects that integrate information across cell lines. DEMETER2 provides absolute gene-level scores and uses a Bayesian inference method for the parameter estimation [11]. The gespeR method uses a regression model to account for sequence-dependent off-target effects, based on the TargetScan model for predicting relationships between siRNA and its off-targets [10]. TargetScan is a miRNA target prediction model that predicts mRNA fold change between wild-type and knockout cells based on various features of miRNA sequence[14].

Discovering cancer dependencies, including synthetic lethality and self-dependencies, is of crucial significance for identifying new cancer treatment options. Synthetic lethality refers to the interaction of two genes, where the simultaneous loss of function through either genetic events or inhibition results in cell death, but the loss of function of either gene alone does not. Several computational methods have been previously proposed for identifying synthetic lethality interactions using loss-of-function RNAi and CRISPR screens [15-19]. Cancer self-dependency refers to a dependency type in which the loss of function of a specific gene leads to cell death preferentially in cells with specific molecular characteristics, such as mutations in the same gene. Studies have previously examined self-dependency in relation to missense and damaging mutations, as well as copy-number amplification, and have identified novel putative cancer genes through these investigations [20-22].

In this research, we investigated the potential of using shRNA-level viability scores instead of gene-level data to enhance the statistical power for identifying cancer self-dependencies. To tackle the challenge of off-target effects, we developed a statistical method for analyzing dependency scores from perturbation screens at the shRNA level. Our hypothesis was that analyzing at the shRNA level would result in greater statistical power for uncovering hidden cancer dependencies. To evaluate the performance of our proposed method, we analyzed pan-cancer and cancer-specific analyses on the Project DRIVE dataset, an RNAi screening project that used deep coverage shRNA lentiviral libraries to target genes across 398 cell lines, providing a valuable resource for exploring and discovering cancer dependencies. In addition, we conducted simulation studies to evaluate the type-I error and statistical power of our approach across various sample sizes.

## MATERIALS AND METHODS

### shRNA viability data from Project DRIVE

In this research, we used the Project DRIVE data to find novel cancer dependencies. Project DRIVE conducted knockdown experiments in three different pools namely poolA, poolB, and BGPD using 158114 shRNAs on 9850 genes in 398 cell lines across 39 cancer types by using a median of 20 pooled shRNA per gene. Pools poolA, poolB, and BGPD included 3492, 3577, and 4178 genes, respectively (Supplementary Figure S1.b). The shRNA viability scores were defined using next-generation sequencing counts as log fold change of shRNA read counts 14 days after the onset of the screen compared to shRNA initial abundance in the input library. For cancer-specific analyses, we only considered 26 cancer types that included at least four cell lines, following completion of all preprocessing steps.

### Gene-level viability data from Project DRIVE

We used ATARiS and DEMETER2 methods for aggregating shRNA viability scores into gene-level dependency scores. ATARiS uses a subset of shRNAs with a consistent pattern of viability scores across cell lines and provides gene-level dependency scores relative to screened cell lines due to using median-centered shRNA viability scores. The ATARiS scores of Project DRIVE are available for 6557 genes in 398 cell lines. On the contrary, DEMETER2 scores are absolute dependency scores computed using a hierarchical Bayesian inference scheme by explicitly modeling the off-target effects associated with the seed sequence of each shRNA. The DEMETER2 scores of Project DRIVE are available for 7975 genes in 397 cell lines.

### Copy number and mutation data

We downloaded the mutation data from the DepMap website and the copy number GISTIC2 data from the CBioPortal website [23]. We considered three different classes of genetic alterations: non-missense mutations, missense mutations, and copy-number amplification. A gene has a non-missense mutation in a cell line if it harbors any of start codon deletion, stop codon deletion, start codon insertion, start codon insertion, frameshift deletion, frameshift insertion, in-frame deletion, in-frame insertion, nonsense mutation, and splice site. Missense mutations are simply defined as mutations annotated with missense mutation. We used the GISTIC2 to determine the copy-number status of a gene. We specifically used GISTIC2 values 2 and -2 representing copy number amplification and deep deletion, respectively. Finally, a gene is considered wild-type in a cell line if it does not harbor missense, non-missense, and copy-number amplification and deep deletion.

### TargetScan data

We used thermodynamic stability of seed pairing in shRNAs in RNAi screening to incorporate the off-target effects of seed sequences. The thermodynamic stability values of 7-mer seeds of the DRIVE data were extracted from TargetScan [24]. The total number of seeds in the TargetScan data and Project DRIVE are 16384 (4^7^) and 13071, respectively.

### IntOGen data

We used the IntOGen data (version 2020.02.01) for validation of our method. It consists of 568 genes involved in cancer among which 422 and 474 genes were also available in the ATARiS and DEMETER2 scores, respectively [25].

### NBDep algorithm

In this section, we explain our proposed framework for identification of cancer self-dependencies, i.e. observing reduced viabilities preferentially in mutated cell lines in the respective gene upon knocking down (Figure 1.). In particular, we identified genes that indicate dependencies in presence of three classes of genetic alterations namely missense mutation, non-missense mutation, copy number amplification as previously studied in [22]. The NBDep algorithm is as follows:

**Figure 1.**
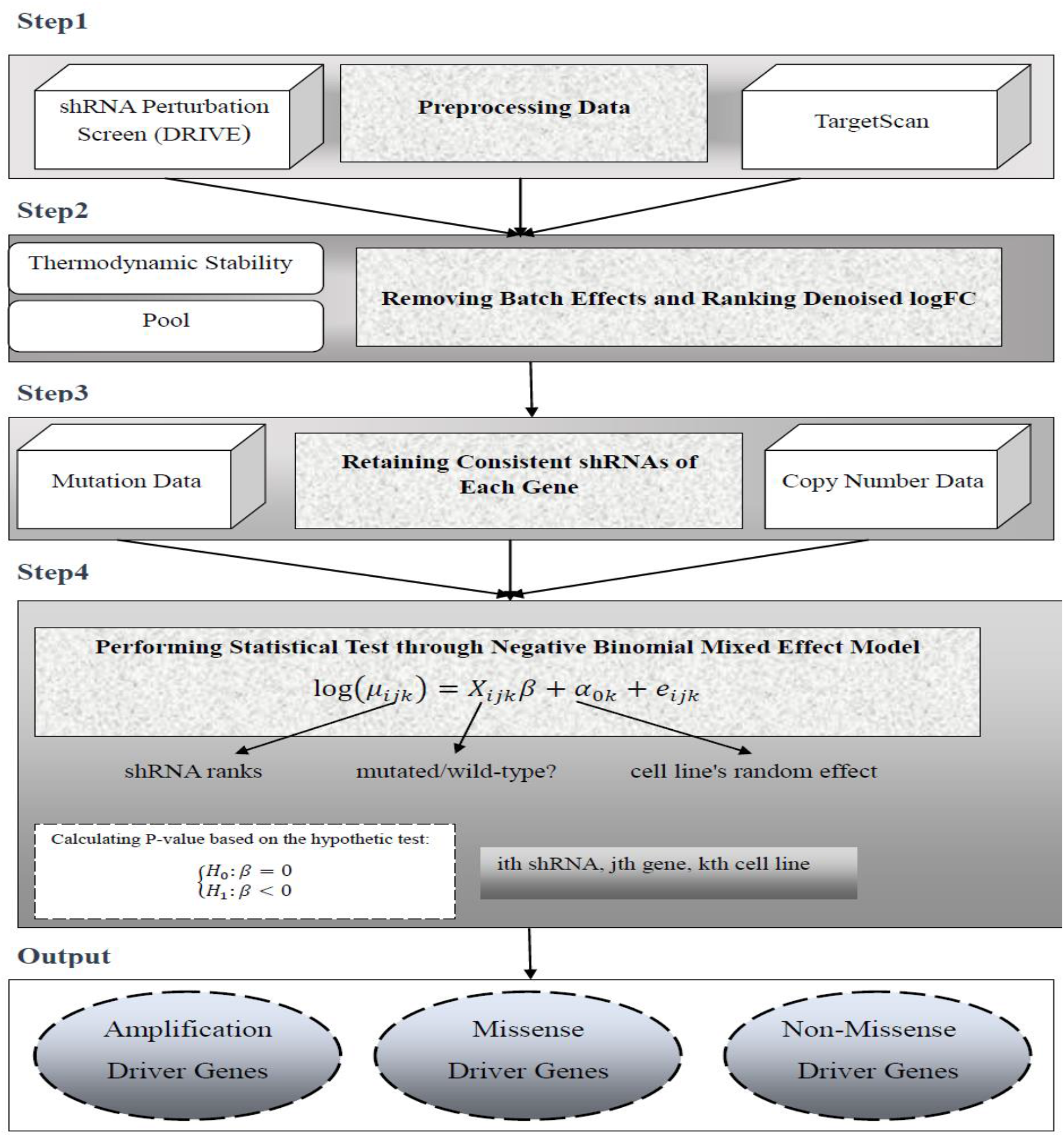
The flowchart of the NBDep algorithm. NBDep employs Project DRIVE, CCLE, GISTIC, and thermodynamic of shRNA’s seeds obtained from TargetScan to identify missense, non-missense, and amplification driver genes. After eliminating batch effects arising from shRNA thermodynamics and pooling, all denoised logFCs in each cell line were ranked. Then, 50% of the most consistent shRNAs of each gene were selected based on their Pearson correlation with the average shRNA profile. Finally, a negative binomial mixed effects model was applied to the shRNA ranks in three alteration classes: missense, non-missense, and amplification alterations. The random effect in this model is associated with the cell line and the fixed effect is a binary variable indicating mutational status.

#### Step 1: Preprocessing

We used the following preprocessing steps to select a subset of shRNAs, genes, and cell lines for further analysis:

1. We eliminated hypermutated cell lines, which were defined as cell lines with a mutation burden three standard deviations above the mean mutation burden across all cell lines.
2. We excluded genes without mutation or copy number data.
3. We eliminated shRNAs from the analysis if the initial abundance level was missing or below 50.
4. In some cases, multiple read counts were reported for a shRNA in a cell line. In these cases, we only retained the average abundance value for the cell line.

The above preprocessing steps resulted in data on 139,407 shRNAs targeting 7,324 genes in 339 cell lines.

#### Step 2: Batch-effect removal and ranking

We then removed two batch effects namely pool and thermodynamic stability of seed sequences from the shRNA data using a linear regression model implemented in the *limma* package in R [26]. Subsequently, we ranked all the corrected shRNA values per each cell line such that shRNA with lower ranks represent higher depletions.

#### Step 3: Obtaining consistent shRNAs

Similar to the ATARiS algorithm, though with a different approach, we found a subset of shRNAs for each gene which exhibited consistent behavior across all cell lines. To achieve this, we calculated Pearson correlation coefficients between all shRNA viability ranks of a gene and the average viability ranks across all shRNAs for all cell lines in a given gene and retained half of the shRNAs with the highest Pearson correlation coefficients. The final preprocessed data to perform NBDep algorithm contains information on 76495 shRNAs targeting 7241 genes 339 cell lines.

#### Step 4: Statistical testing using a Negative Binomial mixed effects Model

Finally, a negative binomial mixed effects model was performed to discover gene drivers in the refined data obtained in the previous steps. To incorporate the heterogeneity of cell lines into the model, they were entered as a random effect covariate in the model. In this model, we assumed that the viability rank for *i*th shRNA and *k*th cell line, denoted by *R*_*ik*_, follows a negative binomial distribution

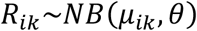

for *i* = 1,2, …, *n*; *k* = 1,2, …, *l*. Parameters *μ*_*ik*_ and *θ* are mean and shape parameter of negative binomial distribution. The probability mass function of *R*_*ik*_ is given by

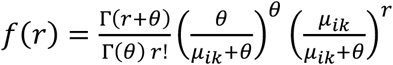

where Γ(.) is the gamma function. The parameter *μ*_*ik*_ is defined as:

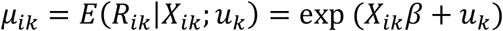

The mutational status of *i*th shRNA in *k*th cell line is a fixed effect, denoted as the variable *X*_*ik*_ where 0 and 1 indicate wild-type and mutant cell lines, respectively. The variable *u*_*k*_ represents the random effect for the *kth* cell line. Under the assumption of this model, the vector *u* = (*u*_1_, *u*_2_, …, *u*_*l*_)^*T*^ follows a multivariate normal with mean 0 and covariance matrix Σ (i.e., *u*∼*MVN*(0, Σ)). The maximum likelihood estimates of the model parameters, (*θ, β*, Σ), were obtained using the *glmmTMB* package in R. To test for the dependency between mutational status and viability score of a gene, we performed one-sided hypothesis testing with *H*_0_: *β* = 0 and *H*_1_: *β* < 0. Under the null hypothesis, *β* follows a normal distribution and consequently the p-value is straightforward to compute.

### Simulation study

In order to evaluate the performance of NBDep and gene-level methods, we conducted two simulation studies to assess type I error and statistical power.

### Simulation study: type I error analysis

In this simulation study, we selected a set of genes for type I error analysis, including known cancer genes *KRAS, NRAS, BRAF, TP53, CDK4, AXIN1*, as well as a few genes that are not known to be involved in cancer namely *UGT8, TRPV3, FUS*, and *BTNL2*. We assumed that there were *m* mutated and *w* wild-type cell lines with respect to a given gene. To generate 5000 random datasets, we performed the following steps:

1. We randomly selected *n*_*m*_ cell lines from the set of all cell lines and assigned them to the mutant group. We generated *n*_*m*_ from a uniform distribution between 2 to *m*.
2. We randomly selected *n*_*w*_ cell lines from the remaining cell lines and assigned them to the wild-type group. We generated *n*_*w*_ from a uniform distribution between 2 to *w*. This ensured that both groups had the same distribution, and the generated data was suitable for assessment of type I error.
3. We then tested for a relationship between mutation status and viability using various methods and calculated type I error as the fraction of rejected tests over the total 5000 tests for each method.

### Simulation study: statistical power analysis

In this section, we examined three known cancer genes: *KRAS, NRAS*, and *PIK3CA*. These genes were highly significant by using NBDep and statistical tests on gene-level data on all available cell lines. These genes were regarded as true positives in this analysis. To analyze statistical power, we simulated datasets in which the distributions of two groups were different. To this end, we subsampled *n*_*m*_ mutated cell lines and *n*_*w*_ wild-type cell lines, where *n*_*m*_ was between 2 and 10, and *n*_*w*_ = *qn*_*m*_ for each gene. The parameter *q* represents the ratio between mutated and wild-type cell lines, and we considered values of *q = 1, 2, 3, 4*. For each gene, *n*_*m*_ and *q*, we generated 800 datasets. The statistical power was then calculated as the ratio of rejected tests to the total number of tests for each setting.

### Statistical testing on gene-level data

We applied the Wilcoxon rank sum test and APSiC, a ranked-based statistical approach based on Irwin-Hall and Bates distributions, to identify self-dependencies from perturbation screens [22]. These tests were conducted on gene-level scores obtained from ATARiS or DEMETER2 dependency scores, which are commonly used for aggregating shRNA viability scores targeting a gene into a single gene-level score. We discovered that the statistical power of the Wilcoxon rank sum test was significantly lower than that of APSiC, and as a result, we only included the results of APSiC in our analyses.

### PPI enrichment

We conducted an enrichment analysis on the set of genes identified through our pan-cancer analysis using data from the STRING database, which contains information on protein-protein interactions. The goal of this analysis was to determine if the identified genes had a greater number of interactions with each other compared to a set of randomly selected genes [27].

### Multiple testing correction

For multiple testing correction, we controlled the false discovery rate using Benjamini and Hochberg approach [28]. The significance level was set to 0.05.

### Data availability

The raw shRNA data and ATARiS scores for Project DRIVE were obtained from the Mendeley Data repository (version 4) at https://data.mendeley.com/datasets/y3ds55n88r, version 4. The DEMETER2 dependency scores from Project DRIVE were retrieved from the DepMap project (version 22Q4). Additionally, molecular profiling data was acquired from the DepMap project (version 21Q4). The CCLE GISTIC copy number alteration data was obtained from the CBioPortal website (Cancer Cell Line Encyclopedia, Novartis/Broad, Nature 2012). The thermodynamic stability of the seed was extracted from supplementary data 5 of the original paper [24].

## Results

### Simulated studies

We first assessed the type I error of our method on synthetic data (see Methods). At the significance level of 5%, the estimated type I error of NBDep was 0.067, slightly higher than the expected 5% error. Next, we compared the statistical power of our method with those of APSiC statistical test on ATARiS and DEMETER2 gene-level dependency scores on three well-known missense driver genes (*KRAS, NRAS*, and *PIK3CA*) using subsampling approaches. To determine whether our method is more robust to low sample size than other methods, we generated 800 datasets for each sample size resulting from various numbers of mutant cell lines and values of q (see Methods). In general, the statistical power of the NBDep algorithm is higher than other methods for small sample sizes with q=1 (Figure 2) and other values of q in the analyzed genes (Supplementary Figure S1).

**Figure 2.**
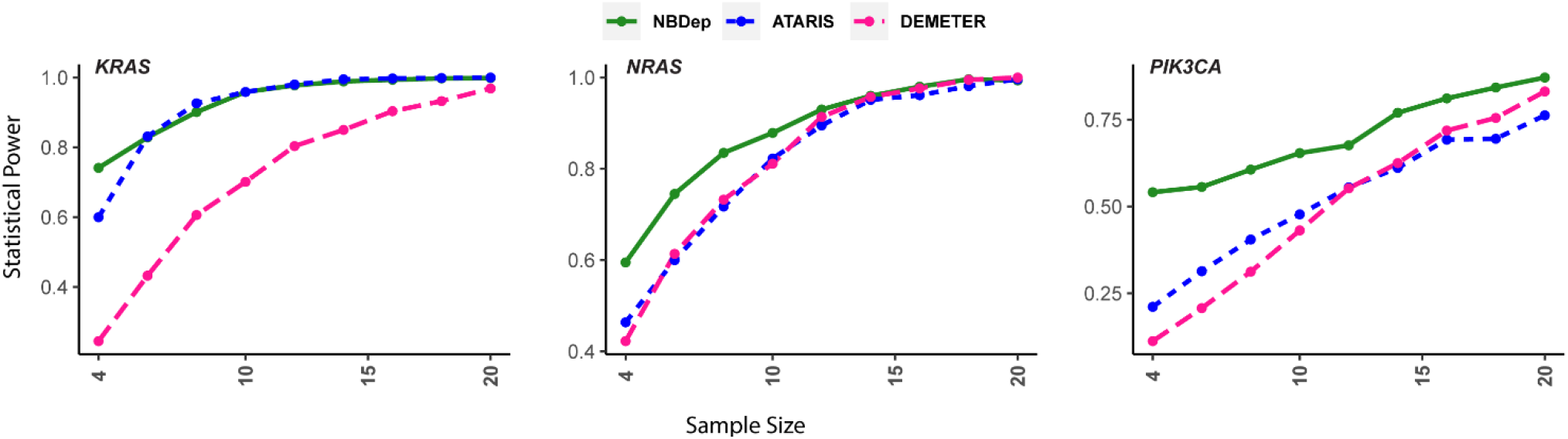
Statistical power of three methods for *KRAS, NRAS*, and *PIK3CA* for various sample sizes and q=1.

### Pan-cancer analysis

NBDep method identified 89 missense mutational driver genes, 41 amplification driver genes, and 125 non-missense mutational driver genes in pan-cancer analyses. In analysis of missense driver genes, NBDep identified sixteen well-recognized genes reported in IntOGen namely *KRAS, NRAS, BRAF, PIK3CA, CTNNB1, TP53, SMAD4, BCL2, SIX1, FBXO11, COL1A1, MAP2K1, TOP2A, DHX9, HRAS* (Figure 3.a). In addition, unreported genes in IntOGen such as *CANT1, IL20RA, BRX1, BUB1, ATIC, Twist2*, and *CSF2RB* were identified by the NBDep method as putative missense mutational cancer genes but these genes were not detected by using APSiC on ATARiS and DEMETER2 gene-level scores. Recently, the important role of these genes as driver genes in different stages of cancer has been presented [29-35]. As depicted in Supplementary Figure S2, there was a significant PPI enrichment observed for both the identified missense driver genes and the identified genes not reported in IntOGen, with p-values of 0.000214 and 0.00767, respectively. One of the top missense driver genes identified by NBDep method is *ANAPC1* which has not been reported in IntOGen and the literature research and it can be introduced as a novel cancer gene. Figure 4.b indicates that *ANAPC1* has significantly lower ranks in missense cell lines than wild-type ones even before taking account for cell line effect in the negative binomial mixed effects model.

**Figure 3.**
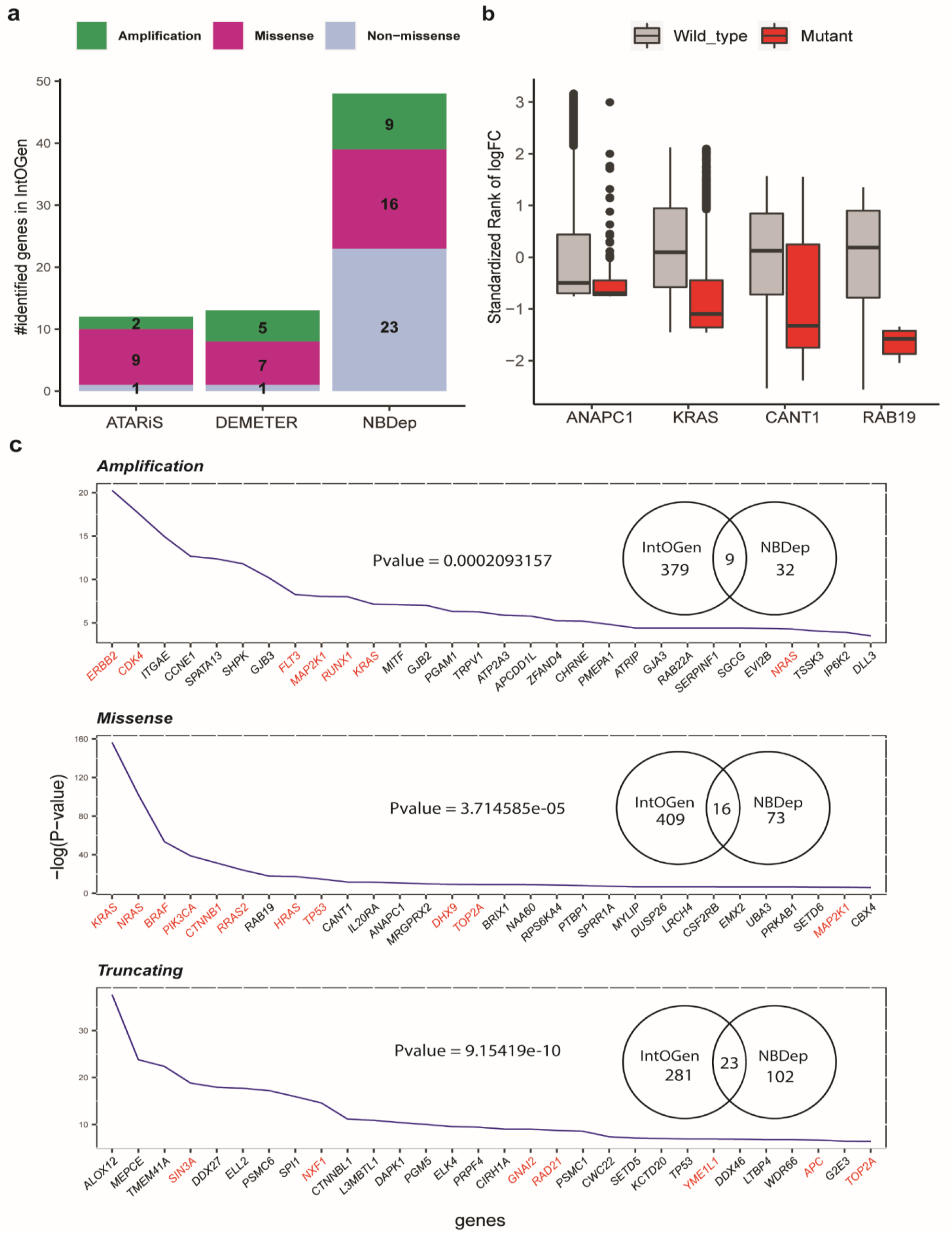
**a**. The number of identified genes common with IntOGen in pan-cancer analysis resulted from three methods: NBDep as well as applying the APSiC method on ATARiS and DEMETER gene-level scores in three alterations: amplification, missense, and non-missense. **b**. Box plots of standardized denoised ranks of one novel missense-driver gene, *ANAPC1*, one experimentally curated missense driver gene, *KRAS*, two genes previously reported in literatures, *RAB19* and *IL20RA*, identifed by NBDep. **c**. Top identified amplificatio, missense mutational, and non-missense genes by NBDep method in pan-cancer. Red color indicates IntOGen genes. Venn diagrams indicates the number of common genes with IntOGen in each of three alteration groups along with their enrichment p-values calculated by the hypergeometric test.

**Figure 4.**
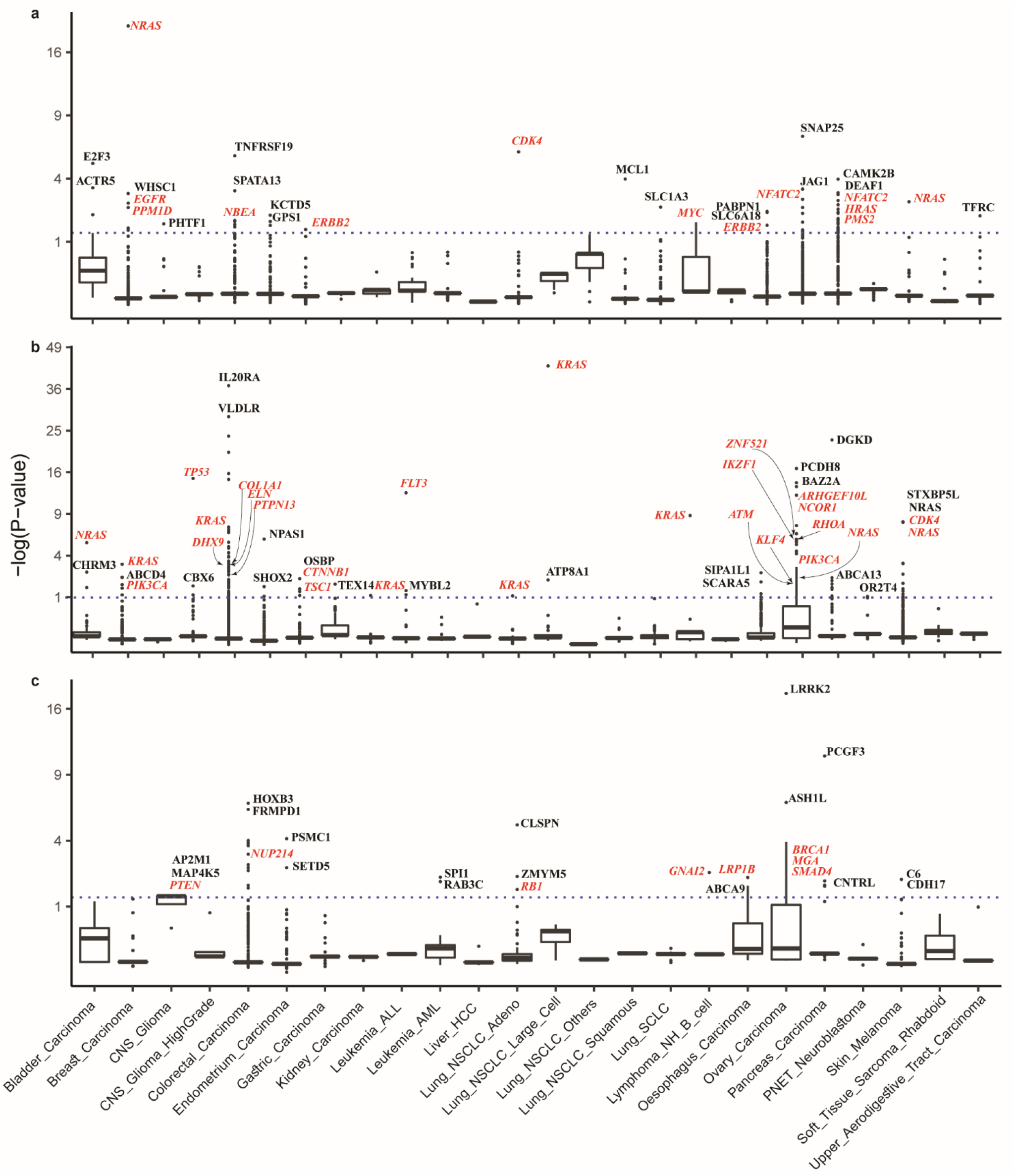
The identified amplification, missense, and non-missense driver genes by NBDep in cancer-specific analyses. The genes highlighted in red are in the IntOGen list.

On the other hand, as depicted in Figure 4a, nine identified amplification driver genes by NBDep are in the IntOGen gene list, namely *ERBB2, CDK4, FLT3, MAP2K1, KRAS, NRAS, GNAS, RUNX1* as oncogenes and *LATS2* as a tumor suppressor. Additionally, other amplification driver genes identified by NBDep, such as *ITGAE, ALKBH3, SHPK*, and *GJB3* have been previously reported to play a significant role in cancer [36-39]. NBDep also identified 23 non-missense driver genes that were previously reported in IntOGen, namely *SIN3A, NTRK1, FN1, MGA, MAP3K1, PTEN, DROSHA, WRN, RELA, CDX2, KDR, CDKN1B, POLD1, PIK3R1, NUP214, PML, GATA3, TOP2A, APC, TP53, RAD21, GNAI2*, and *NXF1*. Recent studies have also highlighted the role of some of the top non-missense driver genes that were not previously identified by IntOGen, such as *ALOX12* [40]. Furthermore, NBDep identified two putative non-missense driver genes *MEPCE* and *TMEM41A* that were not previously known to be cancer driver genes.

In the pan-cancer analysis of all three alterations, several genes were identified as both non-missense and missense driver genes, including *ATIC, MCRS1, TBL3, CACNA1A, TOP2A*, and *TP53*. Moreover, *NRAS, KRAS*, and *MAP2K1* were found to be amplification and missense driver genes. Additionally, *KCNG1* was identified as a driver gene in cancer by NBDep but has not been previously reported in both non-missense and amplification driver genes. Figure 4a shows that the NBDep algorithm identified more common genes across all three alterations compared to using the APSiC method on ATARiS and DEMETER2 scores. Figure 4c displays the top 30 genes identified by the NBDep algorithm in all three alterations, along with their p-values computed using the hypergeometric test for finding the statistical significance of overlaps between identified genes and the IntOGen gene list. It is worth noting that the number of identified driver genes by NBDep was higher in cancer types with more cell lines such as colorectal, skin, and pancreatic carcinoma.

### Cancer-Specific Analysis

We employed the NBDep algorithm to identify driver genes across 26 cancer types only on genes with at least two mutant and two wild-type cell lines. Our analysis revealed 87 amplification, 177 missense, and 45 non-missense driver genes across cancer types. The most frequent identified amplification driver genes belonged to pancreatic and ovarian carcinoma. The identified missense mutational genes were found in colorectal carcinoma and ovarian carcinoma, and non-missense mutational genes were observed in colorectal carcinoma.

Among the amplification driver genes, *NRAS* (in breast carcinoma and skin melanoma), *ERBB2* (in gastric carcinoma, esophageal carcinoma), and *NFATC2* (in ovarian carcinoma and pancreatic carcinoma) were identified in more than one cancer type. Four genes, namely *KRAS* (in breast carcinoma, non-small cell lung cancer (NSCLC), lymphoma multiple myeloma carcinoma, colorectal carcinoma, and leukemia), *NRAS* (in pancreatic carcinoma, bladder carcinoma, primitive neuro-ectodermal tumors(PNET)), *PIK3CA* (in breast carcinoma and ovarian carcinoma), and *ARHGAP31* (in colorectal carcinoma and pancreatic carcinoma) were identified as missense driver genes in more than one cancer type. Notably, the number of identified driver genes by NBDep was higher in cancer types with more cell lines, such as colorectal, skin, and pancreatic carcinoma.

Among the 89 identified missense driver genes in pan-cancer, 21 genes were detected in multiple cancer types, including known cancer genes such as *TP53, KRAS, NRAS*, and *PIK3CA*. Although most of these genes were identified in colorectal cancer, *PIK3CA, TP53* and *CTNNB1* were detected in other cancer types too (breast carcinoma, central nervous system glioma high grade, and gastric carcinoma). *KRAS, IL2ORA, CANT1, NRAS* and *TP53* were the top genes in different cancers. A recent study indicates *IL2ORA* is an important regulator of oncogenic and immune pathways in colorectal carcinoma [29] (Supplementary Table 1).

*NRAS* (breast carcinoma and skin melanoma), *CDK4* (NSCLC, and *ERBB2* (gastric carcinoma and esophageal carcinoma) were identified as the top identified amplification driver genes in different cancer types (Supplementary Table 1).

Also, we found three top genes which have not been reported in IntOGen, TNFRSF19, GJA3, and GJB2, recently found by researchers to have main role in colorectal cancer [41, 42] (Supplementary Table 1).

Our method introduced *SPATA13* a guanine-factor as a novel amplification driver gene in colorectal carcinoma, required for *MMP9* up-regulation via the JNK signaling pathway in colorectal tumor cells. Also, NBDep method identified amplification WHSC1 in breast carcinoma as a novel gene.

Additionally, 16 identified non-missense mutational driver genes in the pan-cancer analyses were also detected as non-missense genes across cancer types, including *CLSPN* and *PIK3C3* (NSCLC), *PSMC1* and *SETD5* (endometrial carcinoma), *MGA* (ovarian carcinoma, *PTEN* (CNS glioma), *SP11* (leukemia), *GNAI2* (lymphoma NH B-cell), and the eight genes in colorectal carcinoma.

For the cancer types with small number of cell lines, the NBDep method is able to identify well-known genes. For leukemia, lymphoma NH B-cell, lymphoma multiple myeloma cancer, and NSCLC that have very low number of cell lines, *KRAS* was detected as the most important missense gene driver. NBDep found *MYC* as an amplification driver gene in lymphoma multiple myeloma cancer while gene-level approches did not identified this gene. Our finding was confirmed by a recent study where amplified *MYC* was shown to be effective in myeloma cancer [43]. In addition, the recent research approves the identified *ATP8A1* by NBDEP as an important gene in NSCLC [44]. The NBDep method also identified 43 missense driver genes in ovarian cancer having a significant protein-protein network (p-value= 2.12e-06) where well-known genes such as *RHOA, PIK3CA*, and *ATM* are among these genes [45-47]. Our method proposed *PCDH8* and *BAZ2A* as novel missense driver genes in ovarian carcinoma.

In summary, the NBDep algorithm is able to identify well-known driver genes in cancer-specific analyses with large and small number of cell lines and to introduce novel putative driver genes (Figure 4).

## Discussion

Perturbation screens including RNAi screening has become increasingly popular in the field of cancer genomics over the past decade. One major limitation of RNAi screening is the off-targets issue where it poses a major challenge to infer actual gene effects in these screens. Different computational methods, including ATARiS and DEMETER2, have been developed to handle off-target effects of shRNAs, leading to gene-level scores known as dependency scores. ATARiS achieves an aggregate score for each gene by discarding shRNAs with non-consistent behavior across all cell lines. DEMETER2 uses a hierarchical Bayesian model to explicitly handle off-target effect associated with seed sequence of each shRNA resulting to gene-level dependency scores.

In this research, we presented a statistical framework aimed at handling off-target effects at shRNA level to identify driver genes associated to missense, amplification, and non-missense alterations. We applied our approach to 26 cancer-specific types as well as pan-cancer data using the Project DRIVE data. We cope with off target effects of shRNAs by incorporating thermodynamic stability of 7-mer seed shRNAs at the shRNA level as a main batch effect proposed by TargetScan. After calculating the denoised shRNA from the original shRNA scores, we then employed 50% of consistent shRNAs of each gene across cell lines. We subsequently performed a negative binomial mixed effects model to investigate association of gene perturbation and alteration statuses of genes. We additionally showed that NBDep algorithm is robust in small sample sizes and can detect driver genes more effectively than using the APSiC method on ATARiS and DEMETER2 gene-level scores.

Having compared the numbers of shRNAs designed for genes in IntOGen to other genes, we recognized that IntOGen genes have more shRNAs than non-well-known genes. This finding advocates that it may be more efficient to design more reagents for all genes when performing RNAi screening.

Our method was capable of identifying well-known genes in pan-cancer and 26 cancer-specific types including *KRAS, NRAS, TP53, BRAF*, and *PIK3CA* in missense alteration, *ERBB2, CDK4*, and *FLT3* as amplification driver genes, and well-recognized non-missense driver genes such as *TP53, TOP2A, APC*, and *GATA3*. Additionally, in cancer-specific analyses, our method was able to identify well-known genes such as *KRAS* and *PIK3CA* as missense driver genes, *NRAS* and *EGFR* as amplification driver genes in breast carcinoma. NBDep method could not identify any driver gene in liver carcinoma. NBDep identified *CCKAR* and *WHSC* as novel missense and amplification driver genes, respectively, in breast carcinoma. *CCKAR* was reported as a driver gene in gallbladder and biliary tract cancer and *WHSC* as *TP53* binding protein [48]. Moreover, *PCDH8*, a protein coding that acts as a cell adhesion molecule, and *BAZ2A* having DNA binding activity, were suggested as missense novel genes in ovarian cancer.

In summary, our method utilizes more information of shRNAs in RNAi screening and is capable to identify above-mentioned self-dependencies in pan-cancer and 26 cancer-specific types while handling off-target effects. Our approach also provides denoised shRNA ranks through which it is possible to explore other types of cancer dependencies such as synthetic lethality.

---

**Supplementary Figure S1.**
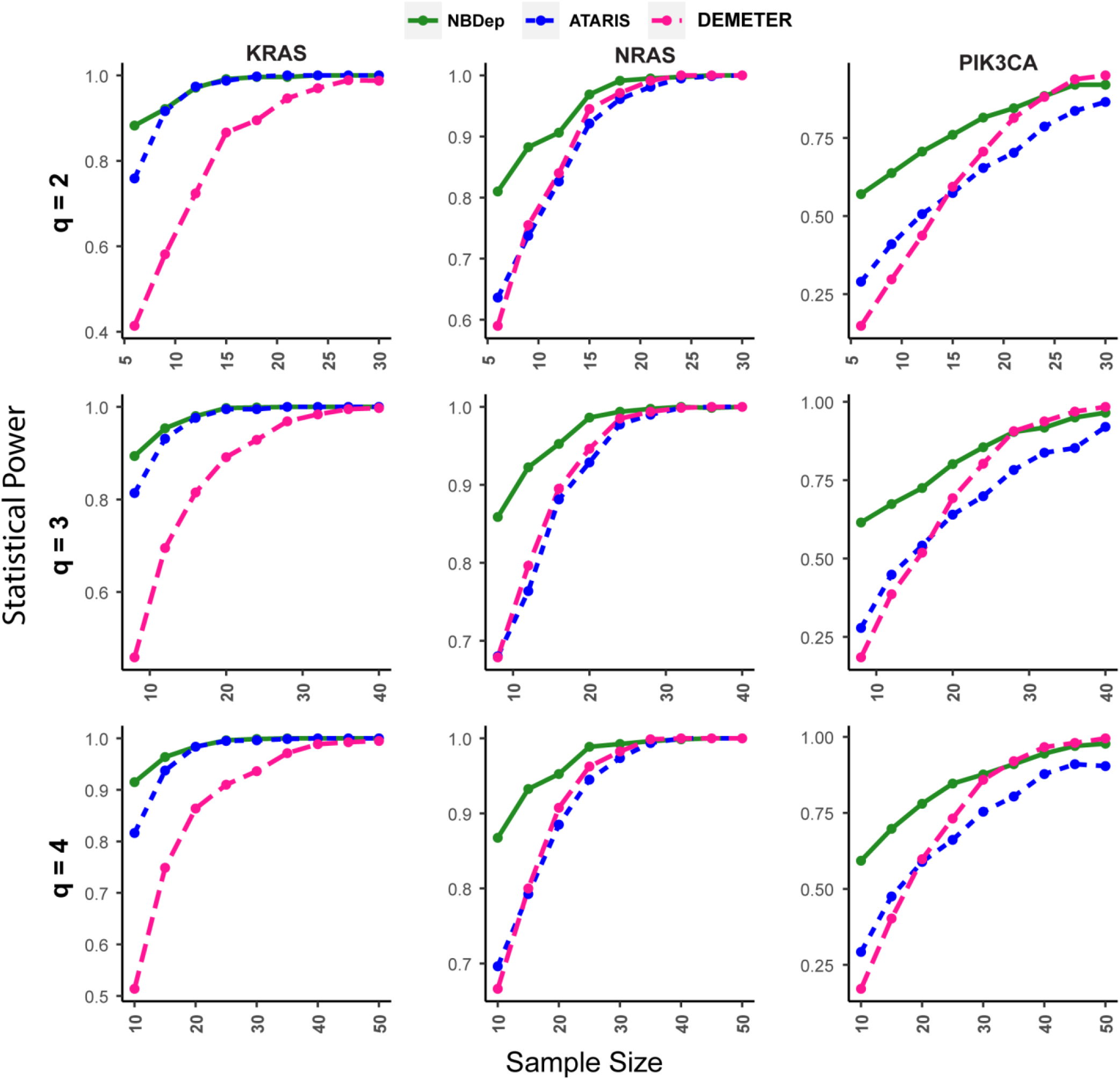
The statistical power of three methods (NBDep, ATARiS, and DEMETER), obtained through a simulation study, for three known driver genes (*KRAS, NRAS*, and PIK3CA) for various sample sizes with q=2,3,4.

**Supplementary Figure S2.**
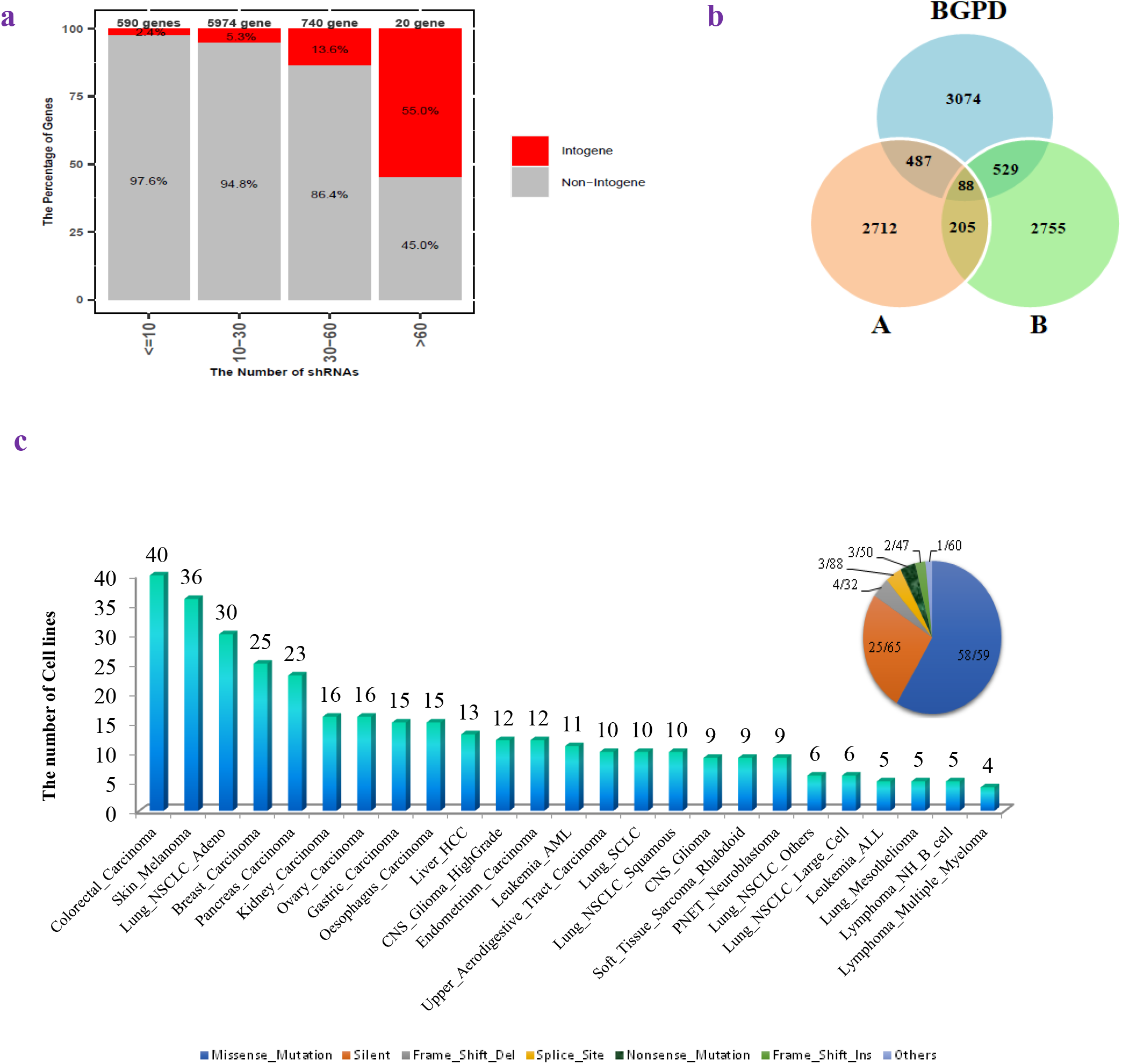
Exploring cell lines, shRNAs, and variant types. **a**. The number of shRNA designed for two groups of genes: IntOGen and Non-IntOGen. **b**. Venn diagram indicating the number of genes in three different pools namely A, B, and BGPD in Project DRIVE. c. The number of cell lines across cancer types in Project DRIVE. The pie chart shows the percentage of each variant type of genes in cell lines, including missense, silent, frameshift deletion, splice site, nonsense, frameshift insertion, and others.

**Supplementary Figure S3.**
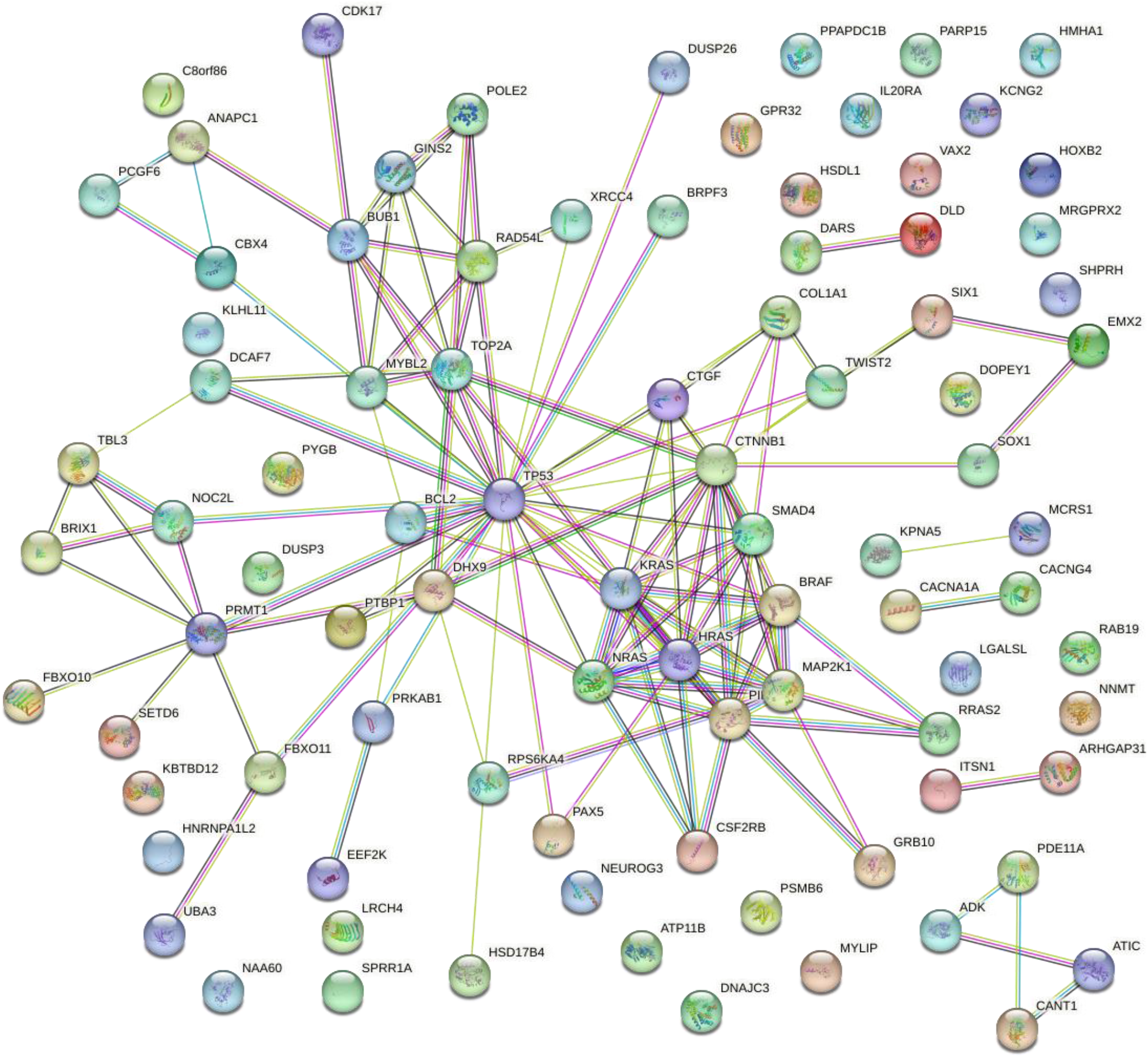
The Protein-Protein network (PPI) of identified missense genes by NBDep in pan-cancer, extracted from STRING. Top identified genes by NBDep including *KRAS, NRAS, PIK3CA, BRAF, HRAS, TP53, TOP2A*, and *CTNNB1* are densely connected in the PPI network. *ANAPC1*, a novel gene identified by NBDep, has coexpressed with *BUB1* which is coexpressed with *TP53* and *TOP2A*, two known cancer driver genes.

**Supplementary Figure S4.**
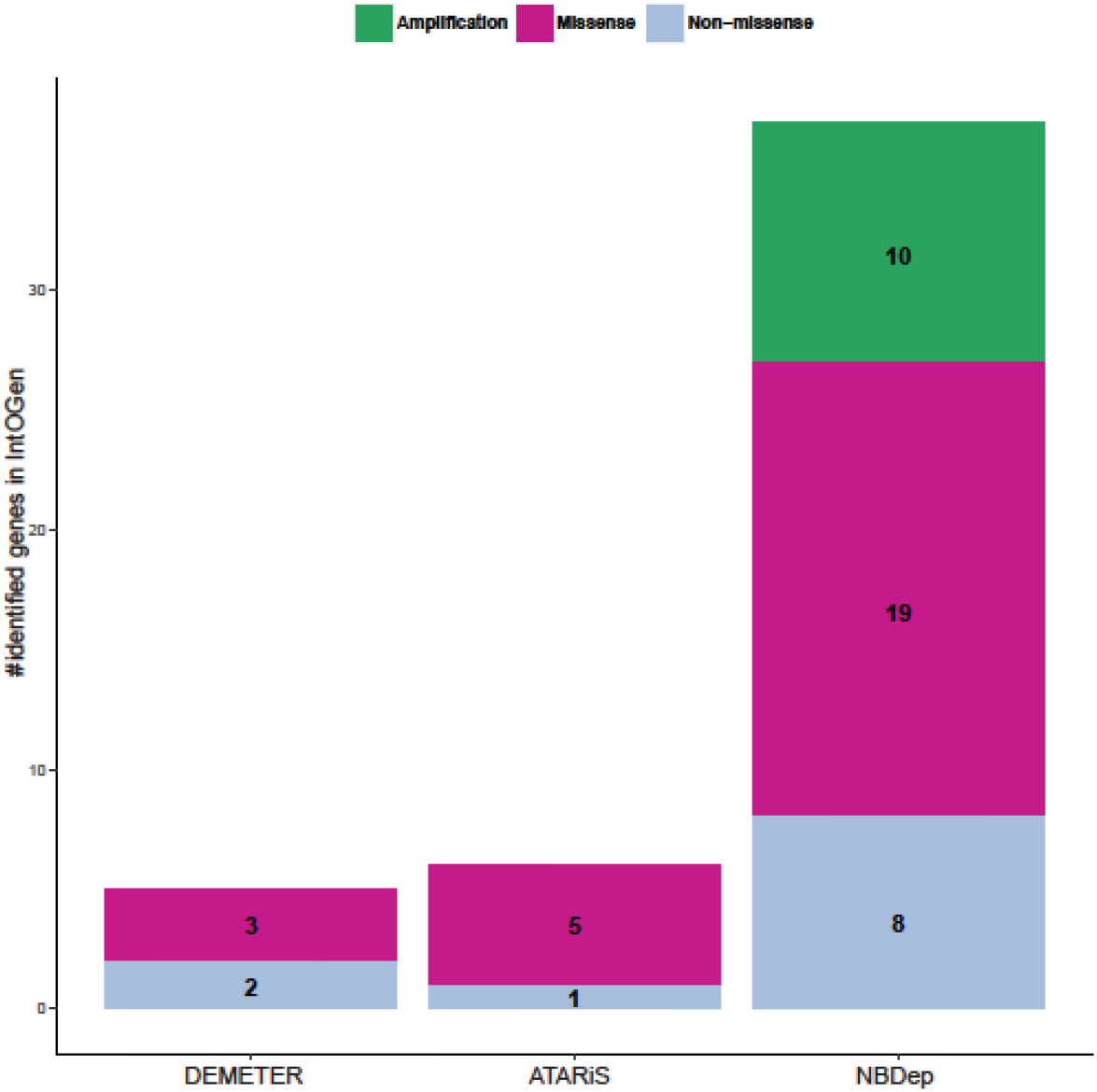
Number of genes identified by three methods (ATARiS, DEMETER, and NBDep) in three alterations (amplification, missense, and non-missense) that are common with IntOGen in any cancer-specific analysis.

**Supplementary Table 1:** Identified genes by the NBDep algorithm in three alterations (amplification, missense, and non-missense) in both pan-cancer and 26 cancer types.

## References

1. Sundara Rajan, et al., Cancer biology functional genomics: From small RNAs to big dreams. Mol Carcinog, 2020. 59(12): p. 1343–1361.

2. Knott Gavin J. and Jennifer A, CRISPER-Cas guides the future of genetic engineering. Science, 2018. 1361: p. 866–869.

3. Dai M., et al., In vivo genome-wide CRISPR screen reveals breast cancer vulnerabilities and synergistic mTOR/Hippo targeted combination therapy. Nat Commun, 2021. 12(1): p. 3055.

4. Arfaoui A., et al., A genome-wide RNAi screen reveals essential therapeutic targets of breast cancer stem cells. EMBO Mol Med, 2019. 11(10): p. e9930.

5. Abdelrahim Maen, et al., RNAi and cancer Implications and applications. Journal of RNAi and gene silencing: an international journal of RNA and gene targeting research, 2006. 2: p. 136.

6. Song C. Q., et al., Genome-Wide CRISPR Screen Identifies Regulators of Mitogen-Activated Protein Kinase as Suppressors of Liver Tumors in Mice. Gastroenterology, 2017. 152(5): p. 1161–1173 e1.

7. Amanda Birmingham and A.K.K. Kozak, RNAi and off-target effects. Frontiers in RNAi, 2014.

8. Brown K. and D. Samarsky, RNAi off-targeting: Light at the end of the tunnel. Journal of RNAi and gene silencing. an international journal of RNA and gene targeting research,, 2006. 2: p. 175–177.

9. Shao D. D., et al., ATARiS: computational quantification of gene suppression phenotypes from multisample RNAi screens. Genome Res, 2013. 23(4): p. 665–78.

10. Schmich F., et al., gespeR: a statistical model for deconvoluting off-target-confounded RNA interference screens. Genome Biol, 2015. 16: p. 220.

11. McFarland J. M., et al., Improved estimation of cancer dependencies from large-scale RNAi screens using model-based normalization and data integration. Nat Commun, 2018. 9(1): p. 4610.

12. Konig R., et al., A probability-based approach for the analysis of large-scale RNAi screens. Nat Methods, 2007. 4(10): p. 847–9.

13. Tsherniak A., et al., Defining a Cancer Dependency Map. Cell, 2017. 170(3): p. 564–576 e16.

14. Agarwal V., et al., Predicting effective microRNA target sites in mammalian mRNAs. Elife, 2015. 4: p. e05005.

15. Shimomura I., Yamamoto Y., and O. T., Synthetic Lethality in Lung Cancer-From the Perspective of Cancer Genomics. Medicines (Basel), 2019. 6(1): p. 38.

16. Dolly S. O., et al., RNAi screen reveals synthetic lethality between cyclin G-associated kinase and FBXW7 by inducing aberrant mitoses. Br J Cancer, 2017. 117(7): p. 954–964.

17. Maia A. F., et al., Genome-wide RNAi screen for synthetic lethal interactions with the C. elegans kinesin-5 homolog BMK-1. Sci Data, 2015. 2: p. 150020.

18. Srivatsa, S., et al., Discovery of synthetic lethal interactions from large-scale pan-cancer perturbation screens. Nature communications, 2022. 13(1): p. 7748.

19. Nigel J. O’Neil, Melanie L Bailey, and P. Hieter, Synthetic lethality and cancer. Nature Reviews Genetics, 2017. 18(10): p. 613--623.

20. Stehr Henning, et al., The structural impact of cancer-associated missense mutations in oncogenes and tumor suppressors. Molecular cancer, 2011. 10: p. 1–10.

21. Kamburov A., et al., Comprehensive assessment of cancer missense mutation clustering in protein structures. Proc Natl Acad Sci U S A, 2015. 112(40): p. E5486–95.

22. Montazeri H., et al., Systematic identification of novel cancer genes through analysis of deep shRNA perturbation screens. Nucleic Acids Res, 2021. 49(15): p. 8488–8504.

23. Mermel Craig H., et al., GISTIC2.0 facilitates sensitive and confident localization of the targets of focal somatic copy-number alteration in human cancers. Genome Biology, 2011. 12(4): p. R41.

24. M., G.D., et al., Weak seed-pairing stability and high target-site abundance decrease the proficiency of lsy-6 and other microRNAs. Nat Struct Mol Biol, 2011. 18(10): p. 1139–46.

25. Martínez-Jiménez, et al., A compendium of mutational cancer driver genes. Nature Reviews Cancer, 2020. 20: p. 555–572.

26. Ritchie, M.E., et al., limma powers differential expression analyses for RNA-sequencing and microarray studies. Nucleic acids research, 2015. 43(7): p. e47–e47.

27. Szklarczyk, D., et al., The STRING database in 2021: customizable protein–protein networks, and functional characterization of user-uploaded gene/measurement sets. Nucleic acids research, 2021. 49(D1): p. D605–D612.

28. Benjamini Yoav and Y. Hochber, Controlling the False Discovery Rate: A Practical and Powerful Approach to Multiple Testing. Journal of the Royal Statistical Society.Series B, 1995. 57: p. 289–300.

29. Yu D., et al., Super-Enhancer Induced IL-20RA Promotes Proliferation/Metastasis and Immune Evasion in Colorectal Cancer. Front Oncol, 2021. 11: p. 724655.

30. Ge J., et al., Expression of biogenesis of ribosomes BRX1 is associated with malignant progression and prognosis in colorectal cancer. Transl Cancer Res, 2020. 9(9): p. 5595–5602.

31. Gerhardt J., et al., The androgen-regulated Calcium-Activated Nucleotidase 1 (CANT1) is commonly overexpressed in prostate cancer and is tumor-biologically relevant in vitro. Am J Pathol, 2011. 178(4): p. 1847–60.

32. Jiang N., Y. Liao, and M.e.a. Wang, BUB1 drives the occurrence and development of bladder cancer by mediating the STAT3 signaling pathway. J Exp Clin Cancer Res, 2021. 40: p. 1–17.

33. Niu N., et al., ATIC facilitates cell growth and migration by upregulating Myc expression in lung adenocarcinoma. Oncol Lett, 2022. 23(4): p. 1–11.

34. Fang X., et al., Twist2 contributes to breast cancer progression by promoting an epithelial-mesenchymal transition and cancer stem-like cell self-renewal. Oncogene, 2011. 30(47): p. 4707–20.

35. Rashid, M., et al., Discovery of a novel potentially transforming somatic mutation in CSF2RB gene in breast cancer. Cancer Med, 2021. 10(22): p. 8138–8150.

36. Tasaki M, et al., ALKBH3, a human AlkB homologue, contributes to cell survival in human non-small-cell lung cancer. British Journal of Cancer., 2011. 104(4): p. 700–706.

37. Franceschi, S., et al., Sedoheptulose Kinase SHPK Expression in Glioblastoma: Emerging Role of the Nonoxidative Pentose Phosphate Pathway in Tumor Proliferation. Int J Mol Sci, 2022. 23(11): p. 5978.

38. Huo Y., et al., GJB3 promotes pancreatic cancer liver metastasis by enhancing the polarization and survival of neutrophil. Front Immunol, 2022. 13: p. 983116.

39. Hu X., et al., ITGAE Defines CD8+ Tumor-Infiltrating Lymphocytes Predicting a better Prognostic Survival in Colorectal Cancer. EBioMedicine, 2018. 35: p. 178–188.

40. Huang Z, et al., ALOX12 inhibition sensitizes breast cancer to chemotherapy via AMPK activation and inhibition of lipid synthesis. Biochem Biophys Res Commun, 2019. 514(1): p. 24–30.

41. Liu Y. J., et al., An Analysis Regarding the Association Between Connexins and Colorectal Cancer (CRC) Tumor Microenvironment. J Inflamm Res, 2022. 15: p. 2461–2476.

42. Schön S., et al., β-catenin regulates NF-κB activity via TNFRSF19 in colorectal cancer cells. Int J Cancer, 2014. 135(8): p. 1800–11.

43. Holien T., et al., MYC amplifications in myeloma cell lines: correlation with MYC-inhibitor efficacy. Oncotarget, 2015. 6(26): p. 22698–705.

44. Li, D., et al., The role of ATP8A1 in non-small cell lung cancer. Int J Clin Exp Pathol, 2017. 10(7): p. 7760–7766.

45. Wang X., et al., Knockdown of RhoA expression alters ovarian cancer biological behavior in vitro and in nude mice. Oncol Rep, 2015. 34(2): p. 891–9.

46. Thorstenson YR, et al., Contributions of ATM mutations to familial breast and ovarian cancer. 63(12), 2003.

47. Campbell I. G., et al., Mutation of the PIK3CA gene in ovarian and breast cancer. Cancer Res, 2004. 64(21): p. 7678–81.

48. Xu H. L., et al., Variants in CCK and CCKAR genes to susceptibility to biliary tract cancers and stones: a population-based study in Shanghai, China. J Gastroenterol Hepatol, 2013. 28(9): p. 1476–81.

